# Kincore: a web resource for structural classification of protein kinases and their inhibitors

**DOI:** 10.1101/2021.02.12.430923

**Authors:** Vivek Modi, Roland L. Dunbrack

## Abstract

The active form of kinases is shared across different family members, as are several commonly observed inactive forms. We previously performed a clustering of the conformation of the activation loop of all protein kinase structures in the Protein Data Bank (PDB) into 8 classes based on the dihedral angles that place the Phe side chain of the DFG motif at the N-terminus of the activation loop. Our clusters are strongly associated with the placement of the activation loop, the C-helix, and other structural elements of kinases. We present Kincore, a web resource providing access to our conformational assignments for kinase structures in the PDB. While other available databases provide conformational states or drug type but not both, KinCore includes the conformational state and the inhibitor type (Type 1, 1.5, 2, 3, allosteric) for each kinase chain. The user can query and browse the database using these attributes or determine the conformational labels of a kinase structure using the web server or a standalone program. The database and labeled structure files can be downloaded from the server. Kincore will help in understanding conformational dynamics of these proteins and guide development of inhibitors targeting specific states. Kincore is available at http://dunbrack.fccc.edu/kincore.

## INTRODUCTION

Protein kinases perform many roles in cell signaling and are widely studied as drug targets with molecules designed to inhibit the active state or various inactive states (1,2). The understanding of conformational dynamics in protein kinases is therefore critical for development of better drugs and novel biological insights. Among the 497 typical kinase domains in the human genome (3,4), currently the structures of 284 have been experimentally determined either in apo form or in complex with ATP or inhibitors. The protein kinase fold consists of an N-terminal lobe, which is formed by five beta sheet strands and one alpha helix called the C-helix, and a C-terminal lobe which consists of five or six alpha helices. The two lobes form a deep cleft in the middle region of the protein creating the ATP-binding active site.

One of the most critical elements in binding ATP and substrate is the activation loop which adopts a unique extended orientation in the active state of the kinase and multiple types of folded conformations in inactive states. It begins with a conserved sequence called the DFGmotif (Asp-Phe-Gly) whose orientation is tightly coupled with the activity status of the protein. The DFGmotif conformations have most often been addressed by using a simple convention of DFGin and DFGout, first defined by Levinson et al. (5). The DFGin group consists of conformations in which DFG-Asp points into the ATP pocket and the DFG-Phe side chain is adjacent to the C-helix. The structures solved in the active state conformation of the enzyme form a subset of this category. In DFGout conformations, the DFG-Asp and DFG-Phe residues swap their positions so that DFG-Asp is removed from the ATP binding site and replaced with DFG-Phe. All Type 2 inhibitors such as imatinib bind to DFGout conformations (6). Some classification systems of kinases are limited to the DFGin/DFGout dichotomy, sometimes with the addition of the position of the C-helix, “in” or “out,” associated with active and inactive kinases respectively (7,8).

The DFGin and DFGout groups, however, provide only a broad description of a more complex conformational landscape (9,10) represented in both crystal structures and studied by molecular dynamics simulations. Other named conformations include “DFGintermediate”, “DFGout-like”, “DFGup”, “pseudo-DFGout,” “auto-inhibited,” and the “CDK/Src-inactive state.” The last of these was first identified in crystal structures of CDK2 (11) and c-SRC (12) and later observed in Abl kinase (5), EGFR (13), IRAK4 (14), and several other kinases. Typically, new structures are classified by comparison to a small number of the earliest solved kinase structures. PDB entries 2CPK (PKA) (15) and 1IR3 (INSR) (16) are used as representatives of active kinases, while 1HCK (11) and 2SRC (12) are representatives of the “CDK/Src-inactive” state. PDB entry 1FGK (FGFR1) (17) is representative of the auto-inhibited form of FGFR and other kinases, and 1IRK (INSR) (18) and 1FPU (Abl1) (19) are the earliest DFGout structures and therefore often treated as canonical. But the criteria for identifying these and other states are not well defined, and the results depend on which structural parameters are used to judge similarity. Even the criteria for judging an active structure or which structures are DFGin and DFGout (20) are not widely agreed upon.

To remedy this situation, in our previous work we developed a scheme for clustering and labeling different conformations of protein kinase structures (21). Our clustering is based on the observation that the first few residues of the activation loop take on a few discrete orientations that can be identified largely based on the position of the side chain of the Phe residue of the DFG motif. We clustered all the conformations into three spatial groups (DFGin, DFGinter, DFGout) based on the proximity of the DFG-Phe side chain to two specific residues in the N-terminal domain. Within these groups, we further clustered the structures by the dihedral angles that determine the location of the DFG-Phe side chain: the backbone ϕ and ψ dihedrals of the X, D and F residues (where X is the residue before the DFGmotif) and the χ_1_ dihedral angle of the Phe side chain. The kinase states are therefore named after the region of the Ramachandran map occupied by the X, D, and F residues (A for alpha, B for beta, L for left-handed) and the Phe χ_1_ rotamer (plus, minus, or trans for the +60°, -60°, or 180° conformations). The importance of the backbone and side-chain dihedrals of the DFGmotif residues has been noted frequently (5,22-26).

In our clustering, among the DFGin structures we distinguished between the catalytically active kinase conformation (labeled BLAminus) and five inactive conformations (in order of decreasing frequency: BLBplus, ABAminus, BLBminus, BLBtrans, BLAplus) (Figure 1). The ABAminus structures strongly resemble the active BLAminus state, save for a peptide flip of the X-D residues, and may therefore be labeled “active-like.” We found in many cases that electron density calculations indicate that they are mismodeled BLAminus conformations (21). The many structures that have been described as the CDK/Src-inactive state mostly fall into the BLBplus state, although some of them are in the closely related BLBtrans state, including the majority of inactive CDK2 structures. The majority of inactive FGFR kinases are in the less common BLAplus state. Among DFGout structures, we identified one dominant conformation labeled BBAminus, which is strongly correlated with Type 2 kinase inhibitors, such as imatinib. Finally, among the small set of DFGinter structures, where the Phe side chain is intermediate between the DFGin and DFGout positions, we distinguished one cluster based on clustering the dihedral angles (BABtrans). The majority of inactive forms of Aurora A kinase are in the DFGinter state

**Figure 1:**
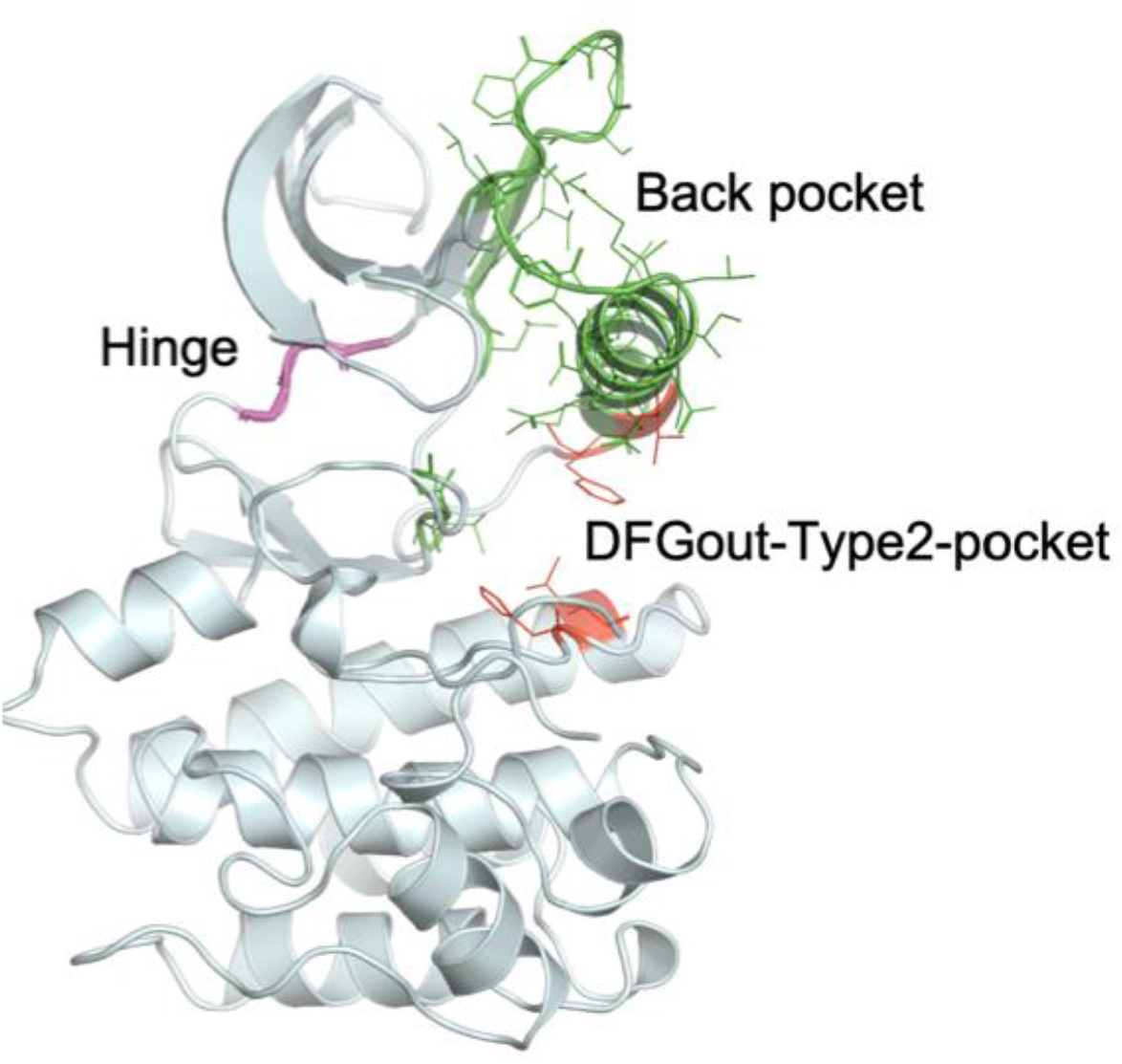
Representative protein kinase structure (3ETA_A) displaying the residues used to define inhibitor binding regions.

Our nomenclature strongly correlates with other structural features associated with active and inactive kinases, such as the positions of the C-helix and the activation loop and the presence or absence of the N-terminal domain salt bridge (21). Since our clustering and nomenclature is based on backbone dihedrals, it is intuitive to structural biologists and easy to apply in a wide variety of experimental and computational studies, as demonstrated recently in identifying the conformation in crystal structure of IRAK3 (27), molecular dynamics simulations of Abl kinase (28), and structural analyses of pseudokinases (29).

In this paper, we present the Kinase Conformation Resource, Kincore – a web resource which automatically collects and curates all protein kinase structures from the Protein Data Bank (PDB) and assigns conformational and inhibitor type labels. The website is designed so that the information for all the structures can be accessed at once using one database table and individual pages for kinase phylogenetic groups, genes, conformational labels, PDBids, ligands and ligand types. The database can be searched using unique identifiers such as PDBid or gene and queried using a combination of attributes such as phylogenetic group, conformational label, and ligand type. We also provide several options to download data – database tables as a tab separated files, and the kinase structures as PyMOL sessions and coordinate files in mmCIF format. The structures have been renumbered by Uniprot and our common numbering scheme, which is derived from our structure-based alignment of all 497 human protein kinase domains (4).

Experimentally determined protein kinase structures in complex with ligands display a diverse array of configurations of the active site. However, examining the conformational dynamics of kinases and its role in ligand binding requires combining two pieces of information – the conformational state of the protein and the type of ligand in complex. Other available resources classify the conformational state (30) or the ligand type (31,32) but not both. Kincore fills a gap by providing a sophisticated scheme for kinase conformations in combination with ligand type labels. We automatically label ligand types based on the pockets to which an inhibitor binds defined by specific residues in the kinase domain. We use five labels for different ligand types as described by Dar and Shokat (33) as further extended by Zuccotto et al. (34): Type 1 – which bind to the ATP binding region only and found in all 8 conformational states; Type 1.5 – ATP binding region and extending into the back pocket (all 8 states); Type 2 – ATP binding region and extending to regions adjacent to the C-helix exposed only in DFGout structures (mostly BBAminus); Type 3 – back pocket only without displacing ATP (predominantly in DFGin-BLBplus and DFGout-BBAminus states); and Allosteric – outside the active site cleft (all DFGin, DFGout, and DFGinter states).

We have also developed a webserver and standalone program which can be used to determine the spatial and dihedral labels for a structure with unknown conformation.

## MATERIAL AND METHODS

### Identifying and renumbering protein kinase structures

The database contains protein kinase domains from *Homo sapiens* and nine model organisms. To identify structures from these organisms the sequence of human Aurora A kinase (residues 125-391) was used to construct a PSSM matrix from three iterations of NCBI PSI-BLAST on the PDB with default cutoff values (35). This PSSM matrix is used as query to run command line PSI-BLAST on the *pdbaa* file from the PISCES server (http://dunbrack.fccc.edu/pisces) (36). *pdbaa* contains the sequence of every chain in every asymmetric unit of the PDB in FASTA format with resolution, R-factors, and SwissProt identifiers (e.g. AURKA_HUMAN). We include only typical protein kinases and exclude “atypical kinases” as defined by UniProt (https://www.uniprot.org/docs/pkinfam).

The structure files are split into individual kinase chains in the asymmetric unit and renumbered according to their UniProt sequences. The necessary alignments are obtained from the Structure Integration with Function, Taxonomy and Sequence (SIFTS) database (37). The SIFTS files were also used to extract mutation, phosphorylation, and missing residue annotations.

The structure files are also renumbered by a common residue numbering scheme using our structure-based protein kinase multiple sequence alignment. Each residue in a kinase domain is renumbered by its column number in the alignment. Therefore, aligned residues across different kinase sequences get the same residue number. For example, in these renumbered structure files the residue number of the DFGmotif across all kinases is 1338 – 1340.

### Assigning conformational labels

Each kinase chain is assigned a spatial group and a dihedral label using our previous clustering scheme as a reference (21). Our clustering scheme has three spatial groups – DFGin, DFGinter, and DFGout. These are sub-divided into dihedral clusters DFGin -- BLAminus, BLAplus, ABAminus, BLBminus, BLBplus, BLBtrans; DFGinter – BABtrans; and DFGout – BBAminus.

To determine the spatial group for each chain, the location of DFG-Phe in the active site is identified using the following criteria:

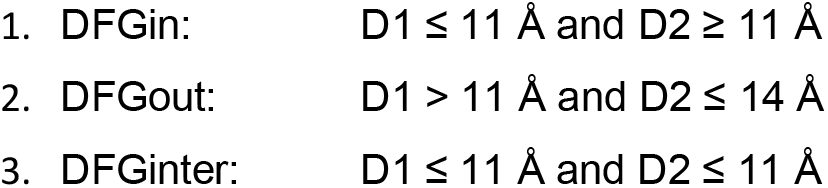

where D1= αC-Glu(+4)-Cα to DFG-Phe-Cζ and D2 = β3-Lys-Cα to DFG-Phe-Cζ. Any structure not satisfying the above criteria is considered an outlier and assigned the spatial label “None.”

To identify the dihedral label the DFG-Phe rotamer type in each chain is identified (minus, plus, trans). The chains for each rotamer type are then represented with a set of six backbone (Φ, Ψ) dihedrals from X-DFG, DFG-Asp, DFG-Phe residues. Using these dihedrals, the distance of each kinase chain is calculated from precomputed cluster centroid points for each cluster with the same rotamer type in the given spatial group. The dihedral angle distance is computed using the following formula,

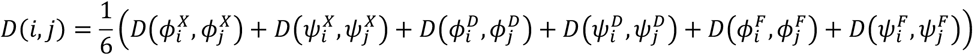

where,

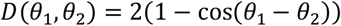

A chain is assigned to a dihedral label if the distance from that cluster centroid is less than 0.45 (about 40°). The chains which have any motif residue missing or are distant from all the cluster centroids are assigned the dihedral label “None.”

The C-helix disposition is determined using the distance between Cβ atoms of B3-Lys and C-helix-Glu(+4). A distance of ≤ 10 Å indicates that the salt bridge between the two residues is present suggesting a C-helix-in conformation. A value of > 10 Å is labeled a C-helix-out conformation.

### Ligand classification

The different regions of the ATP binding pocket used in our analysis (based on sequence numbering of human Aurora A kinase, AURKA_HUMAN) are (Figure 1):

- ATP binding region – hinge residues – residues 211-213
- Back pocket - C-helix and partial regions of B4 and B5 strands, X-DFG and DFG-Asp backbone and DFG-Phe sidechain– residues 166-193, 196-204, 205-207 and 273-275
- Type 2-only pocket – exposed only in DFGout structures – residues 184, 188, 247 and 254

A contact between ligand atoms and protein residues is defined if the distance between any two atoms is ≤ 4.0 Å (hydrogens not included). Based on these contacts we have labeled the ligand types as follows:

1. Allosteric: Any small molecule in the asymmetric unit whose minimum distances from the hinge region and C-helix-Glu(+4) residues are both greater than 6.5 Å.
2. Type 1.5: subdivided as – Type 1.5-front – at least three or more contacts in the back pocket and at least one contact with the N-terminal region of the C-helix. Type 1.5_back - at least three or more contacts in the back pocket but no contact with N-terminal region of the C-helix.
3. Type 2 – three or more contacts in the back pocket and at least one contact in the Type 2-only pocket.
4. Type 3 – minimum distance from the hinge greater than 6 Å and at least three contacts in the back pocket.
5. Type 1 – all the ligands which do not satisfy the above criteria.

### Webserver and software

The program uses the structure file uploaded by the user to extract the sequence of the protein. It aligns the sequence with 10 precomputed HMM profiles of kinase phylogenetic groups. The families came from the UniProt classification (https://www.uniprot.org/docs/pkinfam) with a few modifications (e.g. AURKA is a CAMK kinase) that came from the phylogenetic tree calculated from our structure-based multiple sequence alignment. The 10 HMMs were derived from the structure-based multiple sequence alignment (4) by isolating each family and removing fully-gapped columns. The alignment with the best score among the 10 HMMs is identified and used to determine the positions of the DFGmotif, B3-Lys, C-helix-Glu (+4), and ligand binding residues. The program then computes the distance between specific atoms and dihedrals to identify spatial and dihedral labels and ligand and pocket residues using the assignment method described above.

The standalone program is written in Python3.7. The program is available to download from https://github.com/vivekmodi/Kincore-standalone and can be run in a MacOS, Linux, or Windows machine terminal window. The user can provide individual .pdb or .cif (also compressed .gz) file or a list of files as an input. It identifies the unknown conformation and ligand type from a structure file in the same way as described for the webserver. The scripting and analysis depends on Pandas (https://pandas.pydata.org), and Biopython (38) libraries.

Kincore is developed using Flask web framework (https://flask.palletsprojects.com/en/1.1.x/). The webpages are written in HTML5 and style elements created using Bootstrap v4.5.0 (https://getbootstrap.com/). The 3D visualization is done by using NGL Viewer (http://nglviewer.org/ngl/api/). PyMOL (v2.3) is used for creating download sessions (39). The entire application is deployed on the internet using Apache2 webserver.

## RESULTS

Kincore provides conformational assignments and ligand type labels to protein kinase structures from the PDB. The current update (April 2021) contains structures from 278 kinase genes (284 domains) from humans (7177 chains) and from 57 genes from nine model organisms (884 chains). The PDB files are split by chain, renumbered by Uniprot numbering (37,40) and our common residue numbering scheme, and annotated by conformational and ligand type labels.

The conformational labels are assigned using the structural features and clusters described in our previous work (21). The scheme assigns two types of labels to each chain – a spatial label and a dihedral label. The spatial labels (DFGin, DFGinter, DFGout) are assigned by computing the distance of the DFG-Phe-Cζ atom from the Cα atoms of two conserved residues and assigning a label using distance cutoff criteria (Methods). The dihedral labels within each spatial group are determined by the dihedral angles (φ,ψ of X-DFG, Asp, Phe and χ_1_ for Phe) for each chain which are used to calculate the distance of the structure from the precomputed cluster centroids. A structure is assigned a label if its distance from a centroid satisfies defined cutoff criteria (Methods). All the kinase conformations are represented by a set of eight combined labels. The current statistics of human kinase chains by kinase family and conformational state are given in Table 1.They are: DFGin-BLAminus (active); DFGin-BLBplus (SRC-inactive); DFGin-ABAminus (“active-like”); DFGin-BLBtrans (“CDK-inactive”); DFGin-BLAplus (which we label “FGFR-inactive”); DFGin-BLBminus; DFGout-BBAminus (“type 2 binding”); and DFGinter-BABtrans. The chains that do not satisfy the dihedral distance cutoff criteria for any cluster or are missing some of the relevant coordinates are labeled as ‘None’. Additionally, we have also labeled the C-helix disposition by computing the distance between the C-helix-Glu-Cα atom from the B3-Lys-Cα-atom (as a proxy for the conserved salt bridge interaction) and labeled it as C-helix-in and C-helix-out (Methods).

**Table 1.** Distribution of chains across human protein kinase families and conformational state labels.

| Spatial Label | Dihedral Label | Annotation | AGC | CAMK | CK1 | CMGC | NEK | RGC | STE | TKL | TYR | Other | Total |
| --- | --- | --- | --- | --- | --- | --- | --- | --- | --- | --- | --- | --- | --- |
| DFGin | BLAminus | Active | 306 | 823 | 189 | 1187 | 0 | 0 | 216 | 401 | 545 | 172 | 3839 |
| DFGin | BLBplus | SRC-inactive | 3 | 9 | 0 | 133 | 0 | 0 | 82 | 46 | 358 | 18 | 649 |
| DFGin | ABAminus | Active-like | 60 | 45 | 3 | 95 | 0 | 0 | 11 | 9 | 310 | 88 | 621 |
| DFGin | BLBminus |  | 0 | 7 | 0 | 121 | 1 | 0 | 19 | 18 | 54 | 10 | 230 |
| DFGin | BLAplus | FGFR-inactive | 3 | 2 | 0 | 3 | 0 | 0 | 4 | 40 | 152 | 1 | 205 |
| DFGin | BLBtrans | CDK-inactive | 0 | 0 | 0 | 124 | 0 | 0 | 7 | 1 | 62 | 3 | 197 |
| DFGin | None |  | 14 | 40 | 2 | 76 | 14 | 0 | 26 | 21 | 200 | 24 | 417 |
| DFGinter | BABtrans |  | 0 | 1 | 0 | 0 | 4 | 0 | 2 | 0 | 13 | 0 | 20 |
| DFGinter | None | AURKA-inactive | 10 | 69 | 0 | 15 | 2 | 0 | 7 | 4 | 32 | 0 | 139 |
| DFGout | BBAminus | Type-2 binding | 7 | 6 | 2 | 58 | 0 | 0 | 15 | 96 | 233 | 5 | 422 |
| DFGout | None | DFGout-like | 11 | 10 | 0 | 32 | 9 | 0 | 42 | 14 | 101 | 7 | 226 |
| None | None |  | 7 | 43 | 0 | 87 | 19 | 0 | 17 | 2 | 32 | 5 | 212 |
| Total | - |  | 421 | 1055 | 196 | 1931 | 49 | 0 | 448 | 652 | 2092 | 333 | 7177 |

To assign labels to ligands, we have used specific residue positions to identify regions of the binding pocket – the ATP binding pocket (including the hinge residues), the back pocket, and the Type 2-only region (Figure 1). With the structures renumbered in our common numbering scheme so that all the aligned residues have the same residue number across all the kinases, a ligand is then assigned a label based on its contacts with different binding regions. We have used the following five ligand type labels to annotate all the ligand-bound structures of protein kinases: Type 1 – bind to ATP binding region only; Type 1.5 – bind to ATP binding region and extend into the back pocket (subdivided as Type 1.5-front and Type 1.5-back depending on contact with N-terminal or C-terminal residues of the C-helix, respectively); Type 2 – bind to the ATP binding region and extend into the back pocket and Type 2-only region; Type 3 – bind only in the back pocket without displacing ATP; Allosteric - any pocket outside the ATP-binding region.

In Table 2, we count the number of unique inhibitors in each conformational state and ligand type combination currently in the PDB. The two tables show that Type 1 and Type 1.5 are the most commonly observed inhibitors. However, except for Type 2, all the inhibitor types are observed in complex with multiple conformational states. Both tables demonstrate that Type 1 inhibitors, which only occupy the ATP pocket, bind to truly active structures (BLAminus) only about 60% of the time. ABAminus structures are very similar to BLAminus structures, involving only a peptide flip of the X and D residues of the xDFG motif; the positions of the activation loop and the C-helix are also very similar. The ABAminus state is the second most conformation for Type 1 inhibitors. Type 1.5 inhibitors prefer the back pocket rather than the front pocket between the ATP and C-helix binding sites. Type 3 inhibitors, which occupy the back pocket without competing with ATP, are found in DFGin and DFGout conformations approximately equally.

**Table 2.** Number of unique ligands in each combination of conformation and ligand type.

| Spatial label | Dihedral label | Type 1 | Type 1.5_front | Type 1.5_back | Type 2 | Type 3 | Allosteric | Total |
| --- | --- | --- | --- | --- | --- | --- | --- | --- |
| DFGIN | BLAminus | 1441 | 39 | 106 | - | 9 | 146 | 1686 |
|  | BLBplus | 294 | 6 | 57 | - | 41 | 10 | 399 |
|  | ABAminus | 313 | 6 | 11 | - | - | 12 | 336 |
|  | BLBminus | 142 | 6 | 10 | - | 1 | 6 | 158 |
|  | BLAplus | 53 | - | 52 | - | - | 1 | 105 |
|  | BLBtrans | 135 | 4 | - | - | - | 5 | 142 |
|  | None | 198 | 7 | 22 | - | - | 14 | 235 |
| DFGout | BBAminus | 18 | 1 | 6 | 177 | 36 | 17 | 250 |
|  | None | 44 | 3 | 4 | 65 | 18 | 10 | 144 |
| DFGinter | BABtrans | 7 | - | - | - | - | - | 7 |
|  | None | 57 | 4 | 3 | - | 3 | 3 | 68 |
| None | None | 127 | 5 | 2 | - | 2 | 15 | 149 |
| **Total** |  | 2532 | 80 | 246 | 221 | 109 | 220 | 3295 |

Columns and rows do not add up to the numbers in the “Total” column and “Total” row because some inhibitors bind to more than one conformation and in a small number of cases, the same inhibitor may be found as more than one type because of a single contact.

Many inhibitors are observed in multiple crystal structures bound to one or more different kinases. We counted the number of unique inhibitors that occur bound to kinase chains in two (or more) states across entries in the PDB. In Table 3, we provide the number of unique inhibitors that occur in each pair of states (excluding the unclassified spatial or dihedral labels). The numbers along the diagonal are the counts of unique inhibitors observed in at least one structure of the given state. A total of 242 inhibitors occur in two or more kinase states.

**Table 3.** Counts of inhibitors that are bound to chains in two or more states.

|  | DFGin-BLAminus | DFGin-ABAminus | DFGin-BLBplus | DFGin-BLBminus | DFGin-BLBtrans | DFGin-BLAplus | DFGout-BBAminus | DFGinter-BABtrans |
| --- | --- | --- | --- | --- | --- | --- | --- | --- |
| DFGin-BLAminus | **1686** |  |  |  |  |  |  |  |
| DFGin-ABAminus | 49 | **336** |  |  |  |  |  |  |
| DFGin-BLBplus | 39 | 10 | **399** |  |  |  |  |  |
| DFGin-BLBminus | 25 | 8 | 9 | **158** |  |  |  |  |
| DFGin-BLBtrans | 28 | 3 | 11 | 2 | **142** |  |  |  |
| DFGin-BLAplus | 13 | 5 | 11 | 7 | 1 | **105** |  |  |
| DFGout-BBAminus | 6 | 2 | 3 | 1 | 1 | 2 | **250** |  |
| DFGinter-BABtrans | 5 | 1 | 3 | 3 | 0 | 2 | 0 | **7** |

Numbers along the diagonal (in bold) provide the number of unique inhibitors in each state. The off-diagonal values are the number of unique inhibitors bound to chains in the two states shown in the respective row labels and column headers.

### Website

The web pages on Kincore are designed in a common format across the website to organize the information in a consistent and uniform way. A ‘browse’ page contains all kinase chains in the PDB, with links to all other pages, and a search page allows the generation of pages for specific gene families (AGC, CAMK, CMGC, CK1, NEK, RGC, STE, TKL and TYR (4)), genes (e.g. BRAF), spatial class (DFGin, DFGout, DFGinter), conformational classes (BLAminus, BLAplus, ABAminus, BLBplus, BLBtrans, BLBminus, BBAminus, and BABtrans), ligand types (Type1, Type1.5, Type2, Type3, Allosteric), ligand identifiers (e.g., “DB8”), and individual PDB entries. The page for the gene CDK2 is shown in Figure 2. The table at the top provides counts of each conformation type and representative PDB chains. Each blue field in the table can be clicked to open a page for the respective field (e.g., spatial or dihedral labels or inhibitor name or type). As an advanced query, the database can be queried by simultaneously selecting kinase phylogenetic group, conformational label, and/or ligand type from drop-down menus. For example, a search initiated by selecting TYR group + DFGout-BBAminus + Type 2 ligand type will currently retrieve 193 human chains from 29 genes.

**Figure 2:**
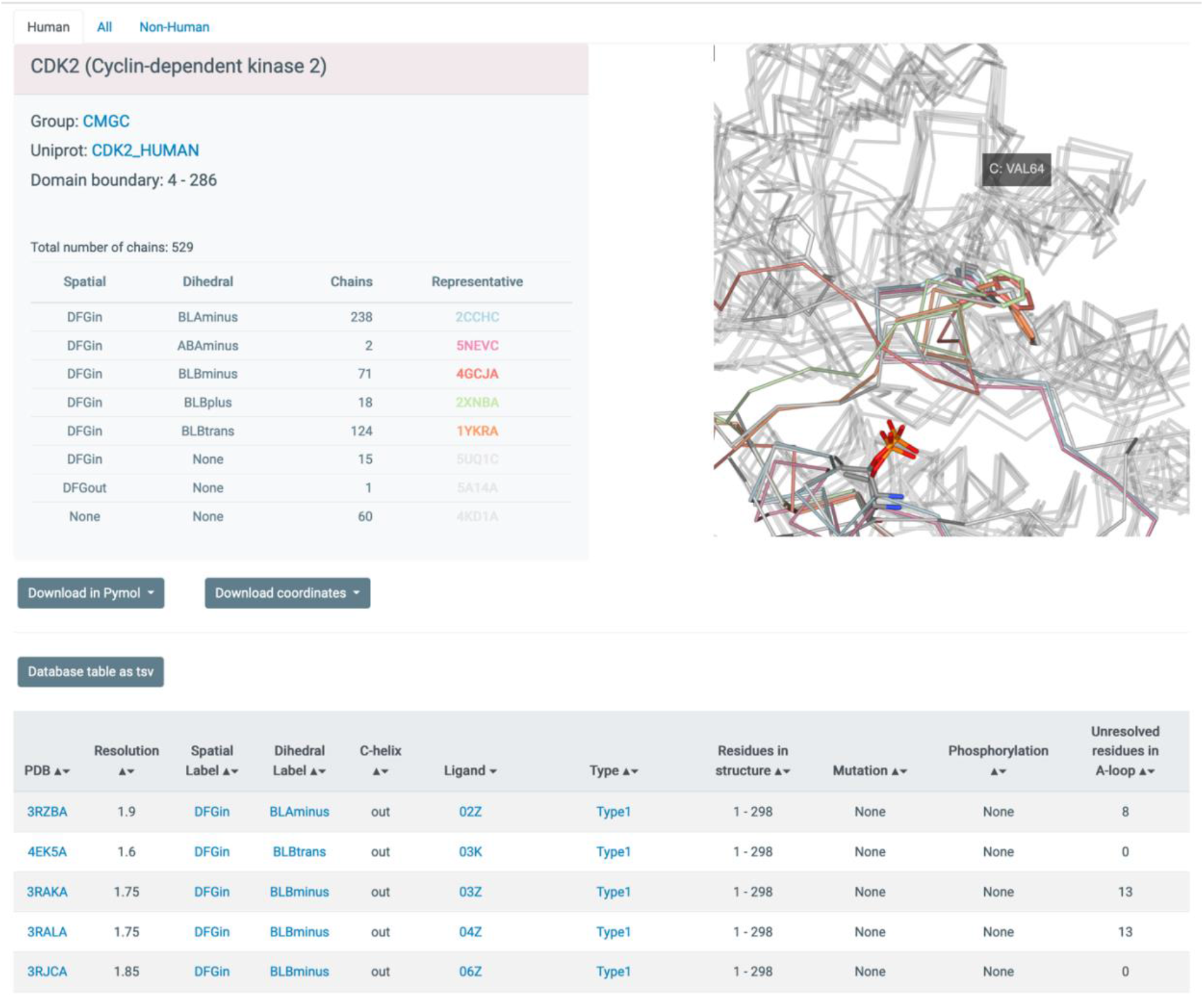
Screenshot of database table displaying entries for PDB chains of CDK2. Every blue string is clickable and will open up a new window with data on all chains corresponding to the link, including the PDB entry (clicking on a chain will lead to a page describing the whole entry), the spatial label (e.g., DFGin), the dihedral label (e.g., BLBminus), the ligand (e.g., 032), and the type of inhibitor (e.g. Type1). Each page provides links to a tab-separated file with the data from the page and to PyMOL sessions and mmCIF coordinate files for all chains listed on the page.

Each page retrieved from the database is organized in two parts – the top part provides a summary of the number of structures in the queried groups or conformations, with representative structures from each category listed and displayed. This is followed by a table from the database with each unique PDB chain as a row providing conformational and ligand type labels and C-helix position, kinase family, gene name, Uniprot ID, ligand PDB ID, and ligand type. The kinase group, gene name, PDB code, conformational labels, ligand name and ligand type are hyperlinked to their specific pages. Each page also contains three tabs at the top to list ‘Human’, ‘Non-human’ and ‘All’ structures. There are buttons provided on each page to download an extended version of the database table as a tab separated file, and to download all of the kinase structures on the page as PyMOL sessions (39) and renumbered coordinate files. In addition to the information displayed on the webpages the tab separated files contain residue number, electron density values (in the form of the EDIA score (41)) for the DFGmotif residues, and R-factor for each chain. Two PyMOL sessions (and scripts in. pml format) are provided for each query – All chains and Representative chains (best resolution, fewest missing residues). Across all the PyMOL sessions, the chains are labeled in a consistent format with their PhyloGroup, Gene, SpatialLabel, DihedralLabel, and PDBidChainid (e.g., TYR_EGFR_DFGin_BLAminus_2GS6A). The coordinate files are provided in mmCIF format with three different numbering systems: the original author residue numbering; renumbered by Uniprot protein sequence; and a common residue numbering scheme derived from our multiple sequence alignment of kinases (4). The same data can also be downloaded in bulk from the ‘Download’ page.

The individual page types provide summary tables and additional pieces of information applicable to each search level. For the Phylogenetic group page, the Summary table provides the number of kinase chains in the group across different conformations with their representative structures (best resolution and least missing residues). These representative structures are also displayed on the page in 3D using NGL viewer. For the Gene page, in addition to the table of chains for a specific gene, the Database table on this page also contains for each chain information on mutations, phosphorylation with total length of the structure and number of residues resolved in the activation loop. The PDB page provides information on each chain the asymmetric unit of a PDB entry, and additionally, the page also contains a sequence feature displaying the UniProt sequence of the protein in the structure and indicating phosphorylated, mutated, and disordered residues. Finally, the ligand page provides access to all chains in complex with a specific ligand. For example, all the structures in complex with ATP can be retrieved by querying for ‘ATP’ on the Search page or clicking on the hyperlinks on the Browse page. This will retrieve 158 chains from 30 genes on the All species tab. Bosutinib (PDB identifier DB8) which is an FDA-approved drug, is found in complex with structures from 10 kinases in 5 different conformations.

The user can upload a kinase structure file in mmCIF or PDB format to determine its conformation and ligand type. We also provide a standalone program written in Python3 which the user can download to assign conformational labels to an unannotated structure and determine the type of ligand in the structure.

### Examples

Kincore enables analysis and visualization of conformational similarity and divergence across sets of structures that result from searches. Because the filenames contain the conformational state (e.g. BLAminus), it is easy to color chains by conformational state. By downloading the coordinate files renumbered by the alignment, it is easy to select the activation loop (residue 1338-1916) and color that accordingly. We provide several examples in Figure 3. PyMol sessions for each image in this figure are provided at http://dunbrack.fccc.edu/kincore/home#comparison_states

**Figure 3.**
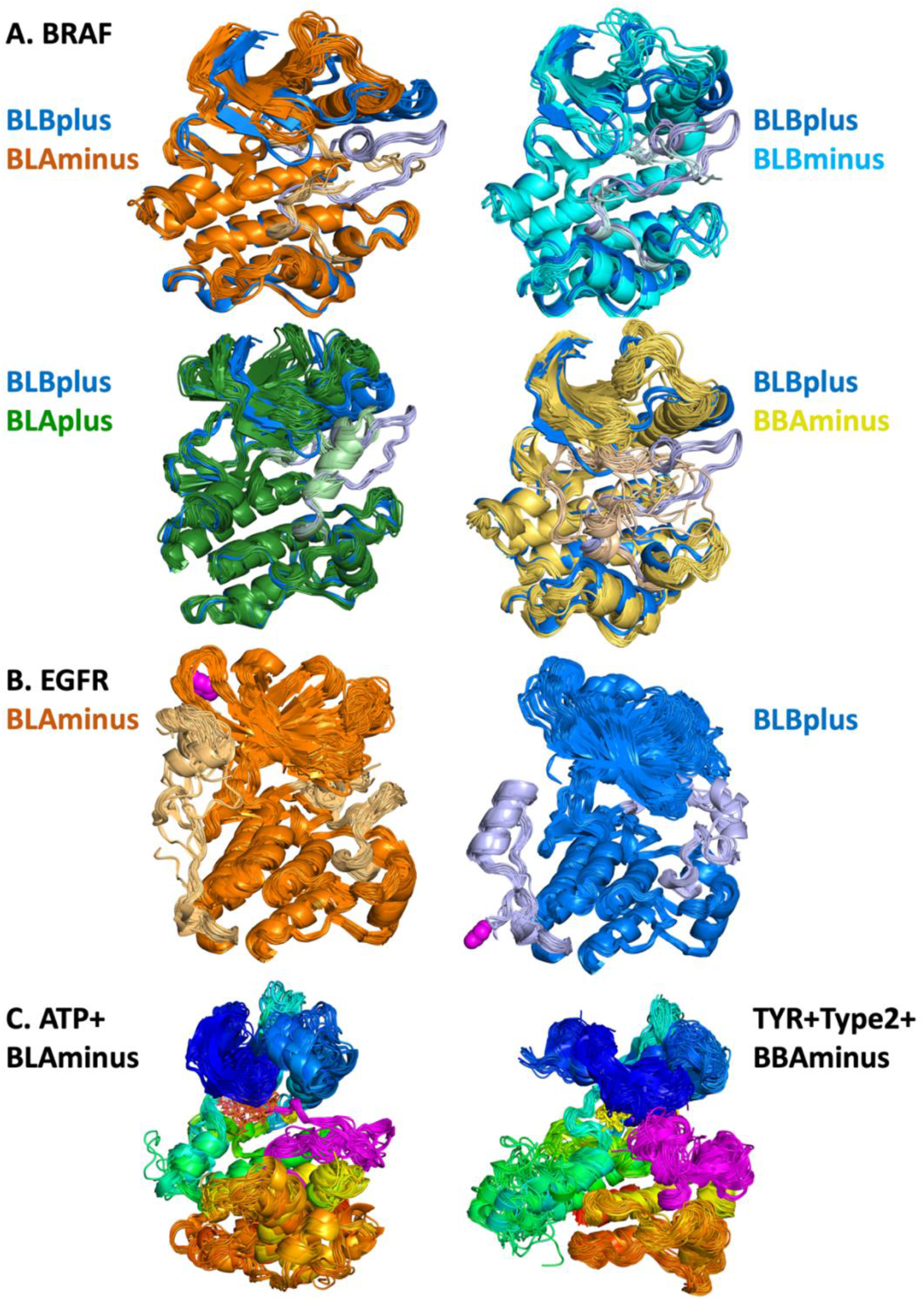
Images of kinase structures that result from searches and downloads from Kincore. **A**. BRAF DFGin-BLBplus structures (“SRC-inactive” conformation in blue with grayish-blue activation loop) compared with DFGin-BLAminus (orange, active conformations), BLBminus (cyan-lightcyan), DFGin-BLAplus (“FGFR-inactive”, forestgreen-lightgreen), and DFGout-BBAminus (Type-2 ligand binding, lightbrown-yellow). A small number of outlier structures in some classes are not shown. **B**. EGFR BLAminus (107 chains, left, orange) and BLBplus (79 chains, right, blue) structures with both the activation loop (right side of each image) and C-terminal tail (left side of each image) shown in lighter colors. The C-terminus of each group is shown in magenta spheres. **C**. ATP-bound BLAminus structures from 14 different kinases in 6 kinase families (left) and BBAminus structures from 29 kinases with bound Type 2 inhibitors (right).

In Figure 3A, pairwise comparisons of 5 different states of human BRAF kinase are shown -- BLBplus (SRC-inactive) with BLAminus (active), BLBminus, BLAplus (FGFR-inactive), and BBAminus (DFGout/Type-2-inhibitor-binding). In each figure, the BLBplus structures are in blue with a grayish blue activation loop. In each case, the position of the activation loop and the C-helix is correlated with the state of the kinase, and differs among the 5 states. As mentioned earlier, we associate the BLAplus state with FGFR kinases (112 out of 127 inactive FGFR1-FGFR4 structures are in this state), although it is found in 20 different kinases, including BRAF. This state was extensively studied by Chen et al. in FGFR1 and FGFR2 (26), who found the change in DFG Phe rotamer from minus in active structures (BLAminus) to plus in PDB 1FGK and 3KY2 (42) (both BLAplus) buried the Phe side chain in a “DFG-latch,” resulting in a highly stable, auto-inhibited structure. A search on Kincore reveals that while one third of the 90 BLAplus conformation of FGFR1 have a fully ordered activation loop, none of the 8 BLBplus structures of FGFR1 are fully ordered

In Figure 3B, the BLAminus and BLBplus states of EGFR are shown in orange and blue with the activation loop on the right side of each figure and the C-terminal tail (residues 971-1020, where the kinase domain is defined as residues 716-970) on the left side of each figure shown in light orange and light grayish blue respectively. The state of the activation loop is highly correlated with the position of the C-terminal tail. In the active BLAminus structures, the tail is mostly coil and reaches up to strands β2 and β3 of the N-terminal domain. In the “SRC-inactive” BLBplus state, the tail contains a helix (residues 993-1002) in contact with the C-terminal domain, and then turns around with the C-terminus in contact with the I-helix of the C-terminal domain. Most of the chains in the BLAminus state are in crystals that contain the asymmetric activating dimer (43), while only two of the 79 BLBplus chains participate in the asymmetric dimer.

In Figure 3C, the results of searches with the Kincore website, including BLAminus+ATP structures and BBAminus structures from the TYR kinase family with bound Type 2 inhibitors. In the BLAminus structures, the position of the DFG Phe (in yellow) and the conformation of the DFGmotif at the beginning of the activation loop (shown in magenta) and the overall position of the activation loop are consistent across the structures.

Finally, we downloaded the predicted structures of human kinases predicted by AlphaFold2 (44) that have been deposited at the European Bioinformatics Institute (http://alphafold.ebi.ac.uk) (45) and ran the Kincore Python3 script to assign our kinase states to the predicted structures. Predicted structures for a total of 496 kinase domains in our list of human kinases were available (skipping PLK5 which is a truncated kinase domain). The distribution of states is compared with structures in the PDB in Table 4 and compared across kinase families in Table 5. Active BLAminus and inactive BLBplus states are more common in the Alphafold predicted structures than in the PDB, among all chains and among chains without ligands (“Apo” in Table 4). DFGout structures are much less well represented in the AlphaFold predictions. Most such structures in the PDB (∼75%) are bound with Type 2 or Type 3 inhibitors. TYR and STE kinases have the most diverse predictions (Table 5).

**Table 4.** Classification of Human Kinase Domain Structures Predicted by AlphaFold2.

| Spatial Label | Dihedral Label | Other names | #AlphaFold2 Chains | AlphaFold2 Percentage | #PDB Chains | PDB Percentage | #PDB Apo Chains | PDB Apo Percentage |
| --- | --- | --- | --- | --- | --- | --- | --- | --- |
| DFGin | BLAminus | Active | 351 | 70.8 | 3839 | 53.5 | 512 | 55.2 |
| DFGin | BLBplus | SRC-inactive | 62 | 12.5 | 649 | 9.0 | 55 | 5.9 |
| DFGin | ABAminus | Active-like | 16 | 3.2 | 621 | 8.7 | 56 | 6.0 |
| DFGin | BLBminus |  | 29 | 5.8 | 230 | 3.2 | 37 | 4.0 |
| DFGin | BLAplus | FGFR-inactive | 6 | 1.2 | 205 | 2.9 | 24 | 2.6 |
| DFGin | BLBtrans | CDK-inactive | 2 | 0.4 | 197 | 2.7 | 12 | 1.3 |
| DFGin | None |  | 8 | 1.6 | 417 | 5.8 | 80 | 8.6 |
| DFGinter | BABtrans |  | 0 | 0.0 | 20 | 0.3 | 5 | 0.5 |
| DFGinter | None | AURKA-inactive | 8 | 1.6 | 139 | 1.9 | 28 | 3.0 |
| DFGout | BBAminus | Type-2 binding | 6 | 1.2 | 422 | 5.9 | 18 | 1.9 |
| DFGout | None | DFGout-like | 5 | 1.0 | 226 | 3.1 | 54 | 5.8 |
| None | None |  | 3 | 0.6 | 212 | 3.0 | 46 | 5.0 |
| Total | - |  | 496 | 100.0 | 7177 | 100.0 | 927 | 100.0 |

The table compares the state of kinase structures predicted by AlphaFold (available from http://alphafold.ebi.ac.uk) and determined with the standalone Kincore Python script with all chains in the PDB and chains without ligands from the Kincore website (http://dunbrack.fccc.edu/kincore/LIGAND/No_ligand)

**Table 5.** Distribution of kinase states by kinase family among 496 AlphaFold2 structure predictions.

| Spatial Label | Dihedral Label | Annotation | AGC | CAMK | CK1 | CMGC | NEK | RGC | STE | TKL | TYR | Other | Total |
| --- | --- | --- | --- | --- | --- | --- | --- | --- | --- | --- | --- | --- | --- |
| DFGin | BLAminus | Active | 55 | 89 | 14 | 53 | 6 | - | 34 | 37 | 50 | 13 | 351 |
| DFGin | BLBplus | SRC-inactive | - | 7 | - | 5 | - | 5 | 13 | 11 | 20 | 1 | 62 |
| DFGin | ABAminus | Active-like | - | 9 | - | - | - | - | - | 1 | 3 | 3 | 16 |
| DFGin | BLBminus |  | 2 | 4 | - | 3 | 7 | - | 9 | 1 | 2 | 1 | 29 |
| DFGin | BLAplus | FGFR-inactive | 2 | - | - | - | - | - | - | 1 | 2 | 1 | 6 |
| DFGin | BLBtrans | CDK-inactive | - | - | - | - | - | - | 1 | - | - | 1 | 2 |
| DFGin | None |  | - | - | - | - | - | - | - | - | 6 | 2 | 8 |
| DFGinter | BABtrans |  | - | - | - | - | - | - | - | - | - | - | - |
| DFGinter | None | AURKA-inactive | - | - | - | - | - | - | - | 1 | 7 | - | 8 |
| DFGout | BBAminus | Type-2 binding | - | - | - | - | - | - | - | - | 6 | - | 6 |
| DFGout | None | DFGout-like | 2 | - | - | - | - | - | 1 | 1 | 1 | - | 5 |
| None | None |  | - | 1 | - | - | - | - | 1 | - | - | 1 | 3 |
| Total | - |  | 61 | 110 | 14 | 61 | 13 | 5 | 59 | 53 | 97 | 23 | 496 |

The table compares the state of kinase structures predicted by AlphaFold (available from http://alphafold.ebi.ac.uk) across different kinase families.

## DISCUSSION

Many proteins exhibit structural diversity across various functional states that can be examined by structural bioinformatics analysis when a large number of experimentally determined structures are available (46). Similar unsupervised learning approaches can be applied to molecular dynamics simulations and other conformational sampling techniques (47). We have previously clustered the conformations of protein kinase activation loops using a dihedral angle metric and a density-based clustering algorithm, DBSCAN. We identified eight conformations of the DFG motif that are correlated with active and inactive functional states, the positions and orientations of other structural elements of the kinase domain including the C-helix and the C-terminal tail, and the types of inhibitors bound to the active site (21). In this paper, we present Kincore, a web-based database that presents our classification of all kinase domains in the Protein Data Bank as well as a standalone program that can be used to classify new kinase structures, molecular dynamics trajectories, or predicted structures of kinases. We also classify the inhibitor type bound to each kinase domain using the standard nomenclature of Type 1 inhibitors (ATP binding site), Type 1.5 inhibitors (binding to the ATP site and a portion of the C-helix pocket), Type 2 inhibitors (binding to the ATP site and the C-helix pocket), Type 3 inhibitors (binding to the C-helix pocket), and allosteric inhibitors binding elsewhere. By combining the classification of the activation loop conformation and inhibitor types, we are able to identify over 200 inhibitors that bind to multiple states of kinases, including 150 that bind to both active and inactive kinases.

Recently, the program AlphaFold2 demonstrated very high accuracy in protein structure prediction from multiple sequence alignments (44). To demonstrate the utility of the Kincore standalone program, we determined the conformational state of 496 human kinase domains predicted by AlphaFold2 available at the EBI (http://alphafold.ebi.ac.uk) (45). AlphaFold2 predicts structures in the active BLAminus conformation for 71% of these kinases, while only 55% of apo structures and 53% of all structures of kinases in the Protein Data Bank are in the BLAminus state. This is not surprising, given that the active conformation must be strongly encoded by the multiple sequence alignment of kinases. We also used ColabFold (48) to generate multiple structure predictions with AlphaFold2 for several individual kinases and ran the Kincore standalone program on these structures. AlphaFold2 was able to generate BLBplus, BLAplus, and BLBminus conformations of some kinases, in addition to the BLAminus states of the same kinases. It is clear that AlphaFold2 can generate different conformational states when run with different random seeds. It will be important in the future to assess how deep learning methods like AlphaFold2 may be used to capture protein dynamics and allostery (49).

Several experimental and computational studies have reported applying the nomenclature from our previous work in structural analyses of kinases Lange and colleagues have solved the crystal structure of the pseudokinase IRAK3 (PDBID 6RUU) and identified its conformation as BLAminus, similar to the active state of a typical protein kinase (27). Paul et.al. have studied the dynamics of ABL kinase by various simulation techniques with Markov state models and analyzed the transition between different metastable states by using our nomenclature (28). Similarly, Maloney et al. have used this classification system to characterize states observed in a molecular dynamics simulation of the V600E mutation of the BRAF kinase (50). Kirubakaran et. al. have identified the catalytically primed structures (BLAminus) from the PDB to create a comparative modeling pipeline for the ligand bound structures of CDK kinases (51). Paul and Srinivasan have done structural analyses of pseudokinases in *Arabidopsis thaliana* and compared with typical protein kinases by applying our conformational labels (29). Finally, Roskoski has performed an extensive comparison of Kincore’s kinase classification of kinase conformations and inhibitor types with his own BRIMR classification of inhibitors (52).

As demonstrated by its utility in analysis of both experimental and predicted structures, we believe that the development of the Kincore database, webserver, and standalone program will greatly benefit a larger research community by making the labeled kinase structures more accessible and facilitating identification of kinase conformations in a wide range of studies.

## AVAILABILITY

The Kincore webserver is available at http://dunbrack.fccc.edu/kincore, and the standalone code for classifying kinase structures is available at https://github.com/vivekmodi/Kincore-standalone.

## ACKNOWLEDGMENT

The authors want to thank Maxim Shapovalov for his help in deploying the server.

